## Extended Data for "Emergent functional behaviors of ribozymes in oxychlorine brines indicate Mars could host a unique niche for molecular evolution"

**Contents:**

**Extended Data Fig. 1** Hammerhead ribozyme and EcoRI enzyme rate constants as a function of perchlorate.

**Extended Data Fig. 2** RNase HII kinetics in perchlorate solutions.

**Extended Data Fig. 3** Broccoli aptamer melting curves in perchlorate solutions.

**Extended Data Fig. 4** TaqI-v2 and HaBlap kinetics and rate constants in perchlorate solutions.

**Extended Data Fig. 5** rPS2.M-heme stoichiometry and Amplex Red oxidation.

**Extended Data Fig. 6** Characterization of chlorination agent and heme turnover for rPS2.M/heme vs. HRP.

**Extended Data Fig. 7** Heme requires rPS2.M to efficiently catalyze chlorination.

**Extended Data Fig. 8** Extinction coefficient of hydrolyzed nitrocefin as a function of salt concentration.

**Extended Data Table 1.** List of oligonucleotides used in this study.

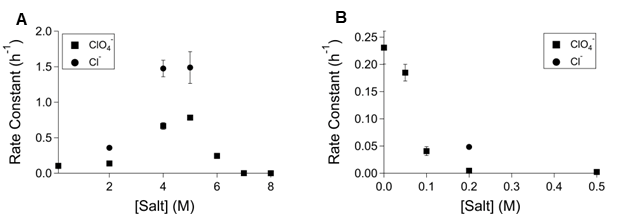

**Extended Data Fig. 1 Hammerhead ribozyme and EcoRI enzyme rate constants as a function of perchlorate.** Rate constant as a function of salt concentration for (**a**) the hammerhead ribozyme (**b**) and EcoRI.

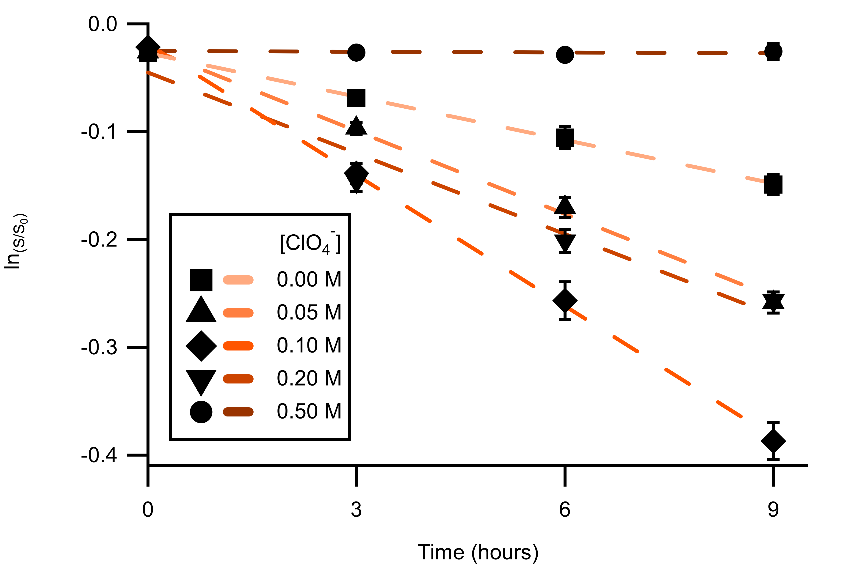

**Extended Data Fig. 2 RNase HII kinetics in perchlorate solutions.**

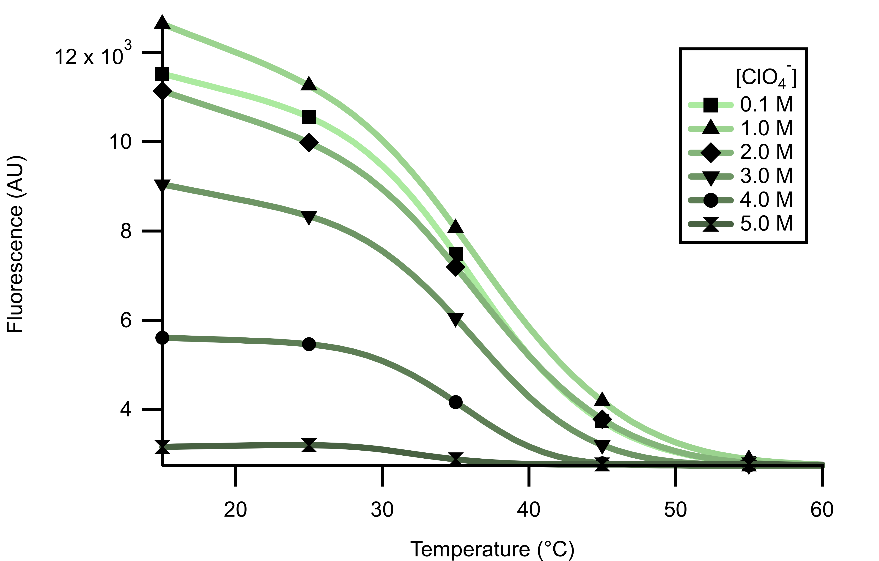

**Extended Data Fig. 3 Broccoli aptamer melting curves in perchlorate solutions.**

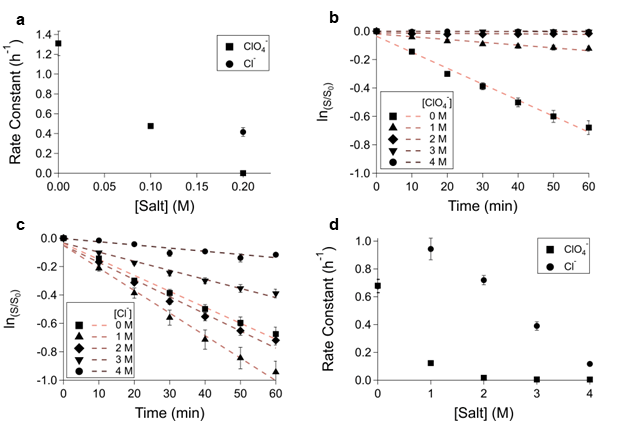

**Extended Data Fig. 4 TaqI-v2 and HaBlap kinetics and rate constants in perchlorate solutions.** **a,** TaqI-v2 rate constant as a function of salt concentration. HaBlap catalyzed nitrocefin hydrolysis in (**b**) perchlorate and (**c**) chloride solutions. **d,** HaBlap rate constant as a function of salt concentration.

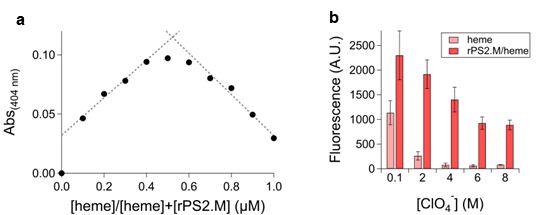

**Extended Data Fig. 5 rPS2.M-heme stoichiometry and Amplex Red oxidation.** **a,** The rPS2.M G quadruplex interacts 1:1 with heme as shown using Job’s method of continuous variation. **b,** The rPS2.M/heme holoenzyme performs peroxidation cycles with hydrogen peroxide and oxidizing Amplex Red to resorufin.

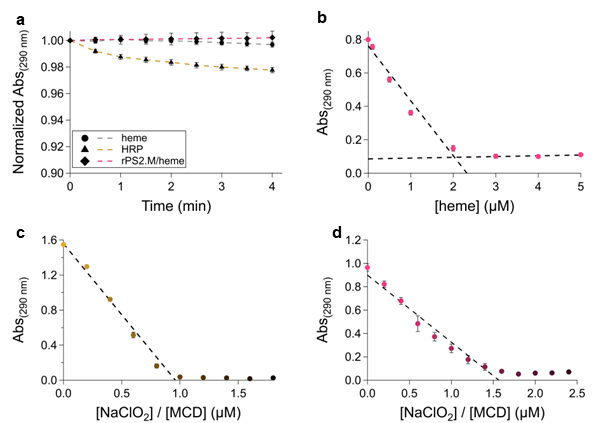

**Extended Data Fig. 6 Characterization of chlorination agent and heme turnover by rPS2.M/heme vs HRP.** **a,** MCD chlorination with heme, rPS2.M/heme, and HRP using sodium hypochlorite instead of sodium chlorite proceeds with HRP, but not heme or rPS2.M/heme. **b,** MCD (50 µM) chlorination with sodium chlorite (200 µM) catalyzed by rPS2.M (7.5 µM) and varying amounts of heme. Based on chlorinating 50 µM MCD with 2 µM heme in limiting heme conditions, and the requirement for two oxidations of heme by chlorite per chlorination event, this gives a turnover number of 12.5 reactions/heme.

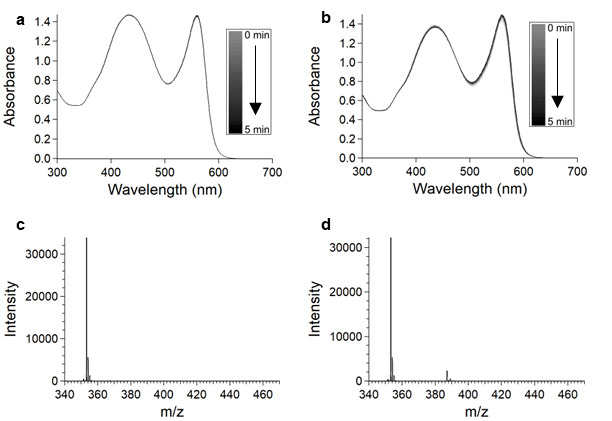

**Extended Data Fig. 7 Heme requires rPS2.M to efficiently catalyze chlorination.** Phenol red chlorination by sodium chlorite in buffer containing no heme (**a**) and buffer containing heme (**b**) monitored by visible light absorbance. Phenol red chlorination by sodium chlorite catalyzed by buffer containing no heme (**c**) and buffer containing heme (**d**) monitored by ESI-MS.

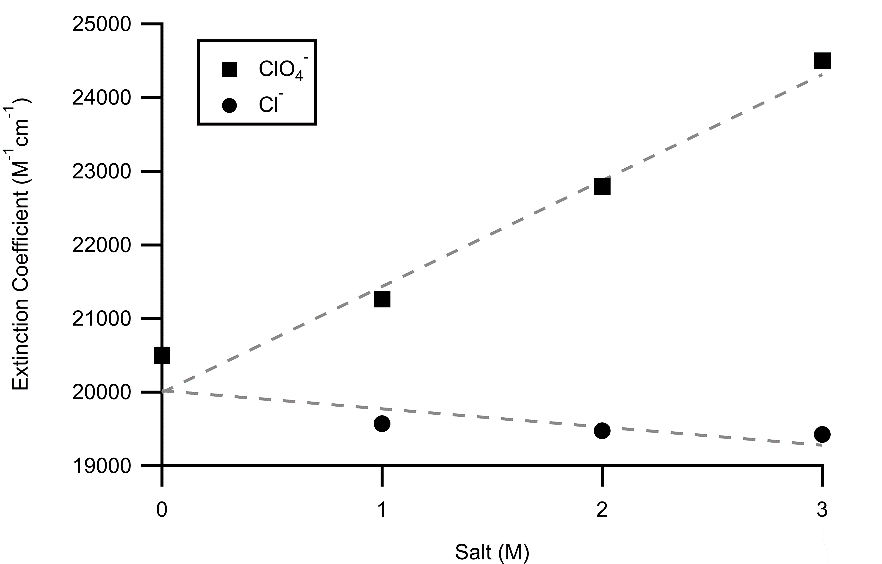

**Extended Data Fig. 8 Extinction coefficient of hydrolyzed nitrocefin as a function of salt concentration.**

**Extended Data Table 1. List of oligonucleotides used in this study.**

| Oligonucleotide | RNA/DNA | Sequence |
| --- | --- | --- |
| EcoRI cleavage assay strand A | DNA | /56-FAM/CGT GTC AGT GAA TTC TTC GAG ATC |
| RNase HII cleavage assay Strand A | DNA/RNA hybrid | /56-FAM/CGT GTC AGT rGAA TTC TTC GAG ATC |
| EcoRI/RNAse HII cleavage assay Strand B | DNA | GAT CTC GAA GAA TTC ACT GAC ACG |
| TaqI-v2 cleavage assay (sense) | DNA | /56-FAM/TCA TAC CTA TTC GAT TGG ATC CTT CCG GTT GGC CGT TAC GGC CTT GCG CTT GAA TTC CGT GCA CAG AA |
| TaqI-v2 cleavage assay (antisense) | DNA | TTC TGT GCA CGG AAT TCA AGC GCA AGG CCG TAA CGG CCA ACC GGA AGG ATC CAA TCG AAT AGG TAT GA |
| Hammerhead ribozyme strand A | RNA | /56-FAM/CGC GCC GAA ACA CCG UGU CUC GAG C |
| Hammerhead ribozyme strand B | RNA | GGC UCG ACU GAU GAG GCG CG |
| Hammerhead displacement strand | DNA | CGC GCC TCA TCA GTC GAG CC |
| Broccoli T7 template (sense) | DNA | **TAA TAC GAC TCA CTA TAG** GAG ACG GTC GGG TCC AGA TAT TCG TAT CTG TCG AGT AGA GTG TGG GCT C |
| Broccoli T7 template (antisense) | DNA | GAG CCC ACA CTC TAC TCG ACA GAT ACG AAT ATC TGG ACC CGA CCG TCT CCT ATA GTG AGT CGT ATT A |
| Broccoli aptamer | RNA | GGA GAC GGU CGG GUC CAG AUA UUC GUA UCU GUC GAG UAG AGU GUG GGC UC |
| tC19Z T7 template (sense) | DNA | **TAA TAC GAC TCA CTA TAG** TCA TTG AAA AAA AAA GAC AAA TCT GCC CTC AGA GCT TGA GAA CAT CTT CGG ATG CAG AGG AGG CAG CCT TCG GTG GCG CGA TAG CGC CAA CGT TCT CAA CAG ACA CCC AAT ACT CCC GCT TCG GCG GGT GGG GAT AAC ACC TGA CGA AAA GGC GAT GTT AGA CAC GCC CAG GTC ATA ATC CCC GGA GCT TCG GCT CC |
| tC19Z T7 template (antisense) | DNA | GUC AUU GAA AAA AAA AGA CAA AUC UGC CCU CAG AGC UUG AG AAC AUC UUC GGA UGC AGA GGA GGC AGC CUU CGG UGG CGC GAU AGC GCC AAC GUU CUC AAC AGA CAC CCA AUA CUC CCG CUU CGG CGG GUG GGG AUA ACA CCU GAC GAA AAG GCG AUG UUA GAC ACG CCC AGG UCA UAA UCC CCG GAG CUU CGG CUC C |
| tC19Z ribozyme |  | GUC AUU GAA AAA AAA AGA CAA AUC UGC CCU CAG AGC UUG AGA ACA UCU UCG GAU GCA GAG GAG GCA GCC UUC GGU GGC GCG AUA GCG CCA ACG UUC UCA ACA GAC ACC CAA UAC UCC CGC UUC GGC GGG UGG GGA UAA CAC CUG ACG AAA AGG CGA U |
| tC19Z Template RNA | RNA | GUC AAU GAC ACG CUU CGC ACG GUU GGC AGA AAA AAA AAA |
| tC19Z RNA primer | RNA | /56-FAM/CTG CCA ACC G |
| tC19Z T7 template for 3 bp hairpin template (sense) | DNA | **TAA TAC GAC TCA CTA TAG** TCA ATG ACT ATT TTT ATA CGG TTG GCA GAA AAA AAA AA |
| tC19Z T7 template for 3 bp hairpin template (antisense) | DNA | TTT TTT TTT TCT GCC AAC CGT ATA AAA ATA GTC ATT GAC TAT AGT GAG TCG TAT TA |
| tC19Z template with 3 bp hairpin RNA | RNA | GUC AAU GAC UAU UUU UAU ACG GUU GGC AGA AAA AAA AAA |
| rPS2.M G quadruplex | RNA | GUG GGU AGG GCG GGU UGG |

T7 promoter sequences are bolded. RNA primer binding sequences for RNA polymerase assays are underlined.
